# Larval motility assay using WMicrotracker™: a high throughput test to discriminate between resistance and susceptibility to anthelmintic drugs in nematodes

**DOI:** 10.1101/2024.05.02.592150

**Authors:** Mélanie Alberich, Marie Garcia, Julie Petermann, Clara Blancfuney, Sophie Jouffroy, Philippe Jacquiet, Anne Lespine

## Abstract

Grazing ruminants suffer from various helminth infections particularly those caused by gastrointestinal nematode (GIN) parasites, which have a considerable impact on their welfare and productivity. Treatment predominantly relies on macrocyclic lactone (ML) anthelmintics, but their widespread application has led to the emergence of drug-resistant parasite populations worldwide. The standard method for detecting resistance, the Faecal Egg Count Reduction Test (FECRT), is susceptible to misinterpretation, leading to flawed management decisions that undermine parasite control efforts. Thus, there is a pressing need for robust resistance detection methods in field parasites. We investigated the potential of the WMicrotracker^TM^ (WMi) motility assay, previously unexplored in ML resistance assessment. The assay first compared ivermectin (IVM) susceptibility among wild-type Bristol N2 (N2B), IVM-selected (IVR10), and *nhr-8* loss-of-function (AE501; *nhr8(ok186)*) *Caenorhabditis elegans* strains. Dose-response curves indicated differences in IVM susceptibility among strains, with IVR10 exhibiting a 2.12-fold decrease in sensitivity compared to N2B. Cross-resistance between IVM, moxidectin (MOX), and eprinomectin (EPR) was explored, demonstrating reduced susceptibility in IVR10 across all drugs compared to N2B. Further investigation was conducted using *Haemonchus contortus* (*H. contortus*) to assess the assay’s applicability in discriminating susceptible from resistant isolates. Results revealed significant differences in drug potency between susceptible and resistant isolates, with MOX demonstrating the highest efficacy. Resistance factors (RF) highlighted the substantial resistance of the resistant isolate to EPR. The motility assay effectively discriminated susceptible from resistant isolates in both *C. elegans* and *H. contortus*. Our findings demonstrate, for the first time, the relevance of the motility assay by WMi as a functional indicator of resistance in nematodes, offering a promising avenue for detecting resistance to MLs. This research sheds light on a novel approach for monitoring drug resistance, vital for effective parasite management strategies.

## 1. Introduction

Parasitic nematodes pose a significant threat to animal health, with profound implications for livestock welfare and productivity. Among them, *Haemonchus contortus* stands out as a highly pathogenic nematode parasite, exacerbating the challenges faced by livestock owners and impacting overall animal well-being and economic output (Fitzpatrick, 2013). Today, the most reliable treatment for these diseases relies on the use of anthelmintic pharmaceuticals. Ivermectin (IVM) belonging to the macrocyclic lactone (ML) class, is the most important anthelmintic drug used worldwide in veterinary and human medicine (Martin et al., 2020). Moxidectin (MOX), another ML was introduced to the veterinary market and has proven to be more effective than IVM against diverse resistant isolates of nematodes in different animal species (Prichard and Geary, 2019). Eprinomectin (EPR) is the only anthelmintic drug that does not necessitate a milk withdrawal period during lactation, ensuring continued accessibility for dairy sheep, goats, and cattle throughout the lactating period. Inevitably, intensive use of an anthelmintic class has selected drug-resistant parasite populations globally in many animal species. This is now a major global problem in small livestock, increasing in cattle (European Medicines Agency, 2016), and in companion animals (Bourguinat et al., 2015). Nowadays, the important losses in productivity in farm animals result from failure to control resistant worms adequately. The rapid spread of MLs resistance compromises not only the control of parasites in animals but also in humans (Doyle et al., 2017; Laing et al., 2017). Consequently, preventing, diagnosing, and managing anthelmintic resistance (AR) take precedence as primary research focuses in veterinary helminthology (Morgan et al., 2019) and is also a concern for control of nematode parasites in humans(Osei-Atweneboana et al., 2011).

In that context, actions are needed to detect drug resistance early, and to preserve efficacy of existing drugs as much as possible. This requires an in-depth understanding of the mechanisms of resistance to anthelmintics. Investigating mechanisms of anthelmintic resistance in parasites is a challenging task because of their complex life cycle, relying on propagation in the host. Therefore, the free-living nematode *Caenorhabditis elegans* is a powerful and recognized model to study anthelmintic resistance (Wit et al., 2022). The use of this model system has considerably improved the understanding of the mechanism of action of anthelmintics, as the targets of each have been elucidated through advanced genetic screens for *C. elegans* mutants that were resistant to their effects (Hahnel et al., 2020). *C. elegans* strains resistant to anthelmintics have been developed and are important models for understanding drug resistance (Driscoll et al., 1989; James and Davey, 2009; Ménez et al., 2016). Moreover, the approach of using *C. elegans* as an experimental model of parasitic nematodes is promising as it allows fast progress. Indeed, thanks to the use of this model, we have recently identified a new key regulator of IVM tolerance; the nuclear hormone receptor NHR-8 (Ménez et al., 2019).

In the field, detecting resistance in gastrointestinal nematode parasites relies on the Faecal Egg Count Reduction Test (FECRT). This test compares egg counts prior to and post a drench treatment, computing a specific drug’s efficacy percentage. However, misunderstanding of potential factors that may affect FECRT results can prompt misguided decisions in management, with significant consequences for continuous parasite control (Morgan et al., 2022). In this context, there is an imperative to develop robust methods for detecting drug resistance in field parasites. Motility tracking system, proposed as a whole animal approach, has been initially developed to fully characterize the locomotor behavior and circadian rhythm of locomotor activity in adult *C. elegans* (Simonetta et al., 2009; Simonetta and Golombek, 2007). Subsequently, the usefulness of this test was quickly understood and multiple applications of the WMicrotracker (WMi) emerged. Indeed, this assay has been used to study dye toxicity in *C. elegans* (Bianchi et al., 2015). High throughput motility analysis has been reviewed for its possible application in the search for new anthelminthics in the context of resistance (Buckingham et al., 2014). As examples, it has been used to screen anthelmintic activity of medicinal plants on *C. elegans* (Liu et al., 2022, 2018) and of essential oil against *H. contortus* (Garbin et al., 2021). The interest in studying various parasites soon became apparent and a number of studies were carried out. It is in that context that WMi helped to demonstrate the promising anthelmintic properties of the repurposing drug EVP4593, as an anthelmintic on L3 of different parasite species such as *Cooperia oncophora*, *Ostertagia ostertagi*, *H. contortus* and *Teladorsagia circumincta* (Liu et al., 2019). Then, the protocol to study worm motility with WMi has been improved as a high throughput test to study anthelmintic activity of large libraries of compounds on *H. contortus* (Taki et al., 2021). In a recent study, Suarez *et al*. investigated the interaction between IVM and EPR using WMicrotracker^TM^ on *C. elegans*. Their findings dissuaded the concurrent usage of these drugs, a practice occasionally suggested by commercial formulations, as the combined effects were not superior to their individual actions (Suárez et al., 2022). Nevertheless, given that the WMi motility assay has not been previously applied in the context of ML resistance, the aim of this research was to evaluate its potential to discriminate susceptible from resistant nematodes. We aimed to assess computer-aided measurements of motility as a method for rapidly evaluating drug efficacy in nematodes and assessing their resistance status. Then, we have evaluated the suitability of such methods for assessing the drug tolerance status to IVM, MOX and EPR of several *C. elegans* strains of known resistance status and tested its applicability on a field parasite of interest, *H. contortus*.

## 2. Materials and methods

### 2.1 Materials

IVM, MOX, EPR, dimethyl sulfoxide (DMSO), bacto agar, bacto peptone, bovine serum albumin (BSA), CaCl2, LB, NaCl and MgSO4 were purchased from Sigma-Aldrich (St Quentin Fallavier, France). For all experiments, IVM, MOX and EPR were dissolved in DMSO.

### 2.2 *C. elegans* nematode strains and cultivation conditions

Wild-type *C. elegans* strain N2 (N2B) and the mutant strain AE501, *nhr-8*(ok186) as well as the OP50 *Escherichia coli* strains were obtained from the *Caenorhabditis* Genetics Center (CGC, University of Minnesota, Minnesota, Minneapolis, MN, USA). IVR10 strain was kindly provided by Dr C. E. James (James and Davey, 2009). All strains were cultured and handled according to the procedures described previously (Ménez et al., 2019, 2016). Briefly, nematodes were cultured at 21°C on Nematode Growth Medium (NGM) agar plates (1.7% bacto agar, 0.2% bacto peptone, 50 mM NaCl, 5 mg/L cholesterol,1 mM CaCl2, 1mM MgSO4, and 25 mM KPO4 Buffer) seeded with *Escherichia coli* strain OP50 as a food source. N2 Bristol and AE501 strains were cultured on classic NGM agar plate while IVM-selected strain (IVR10) was cultured on NGM plates containing 11.4 nM (10 ng/ml) of IVM. IVM-containing NGM plates were prepared as follows: stock solutions of IVM were diluted in NGM at the adequate concentration before pouring plates. Nematodes were synchronized through egg preparation with sodium hypochlorite. Briefly, an asynchronous population with majority of gravid adults and eggs was collected by washing the bottom of the NGM plates with M9 buffer (3 g KH2PO4, 6g Na2HPO4, 5 g NaCl, 0.25 g MgSO4 7H2O in 1 l water) and centrifuged at 1300g for 30 seconds. All larval stages except eggs were lysed with a bleaching mixture (5M NaOH and 1% hypochlorite). Three washes of M9 were done to remove the toxic bleaching mixture. *C. elegans* eggs were then hatched overnight at 21°C, on an orbital shaker, in M9 solution without bacteria to obtain a synchronized population of first-stage larvae (L1).

### 2.3 H. contortus isolates

The *H. contortus* isolate R-EPR1-2022 was originally obtained from a dairy sheep farm in South of France, recently diagnosed with ML-resistance (Jouffroy et al., 2023), while S-H-2022 was a ML-susceptible isolate from the same region, kept in the laboratory since for several generations.. The eggs were isolated from sheep feces collected either from a farm where EPR was effective, or from a farm where EPR resistance had been identified following drug treatment. Since their recovery, isolates were passaged every 2 months in sheep infected with 10000 infective larvae (L3) to maintain the strains. Experimentations involving sheep were performed in the experimental sheepfold facilities of the veterinary school of Toulouse ENVT (accreditation number E31 555 027). The project has been registered and authorized under the number APAFIS #40417-20230119164118 v3 by the French Ministry of Higher Education and Research.

### 2.4 Larval Motility Assay (LMA)

The susceptibility to MLs of the *C. elegans* strains and *H. contortus* isolates was determined in a larval motility assay (LMA). Motility of nematodes was assessed from score activity recording using the WMicroTracker™ One (WMi) from PhylumTech (Santa Fe, Argentina), which detects infrared microbeam interruptions due to worm movement in liquid media. The method used has been adapted from protocols previously described for *C. elegans* (Bianchi et al., 2015; Risi et al., 2019; Simonetta et al., 2009; Simonetta and Golombek, 2007) and *H. contortus* (Munguía et al., 2022). The capacity of IVM, MOX and EPR to inhibit the worm motility was measured in a dose dependent assay by adding the drugs at increasing concentration (0.0125-1 µM for *C. elegans* and 0.01-100 µM for *H. contortus*). The drugs were solubilized in DMSO, and the final concentration of DMSO in the assay was below 0.5 %, to exclude harmful effects of the vehicle. Given the significant procedural differences between *C. elegans* and *H. contortus*, in order to enhance clarity, a schematic overview of the LMA experimental procedure as well as the appropriate timeline inspired from Preez *et al* (Preez et al., 2020), is provided in Fig. 3 for *C. elegans* and in Fig 4. for *H. contortus*.

#### 2.4.1 LMA on *C. elegans* strains

The LMA on *C. elegans* measures the potency of anthelmintics in inhibiting motility in young adults. After bleaching, 4500 L1 were poured on 3 classic NGM agar plates. Synchronized young adults (40-50 per well) were seeded into a final volume of 200 µl M9 in a 96-flat well plate. Plates were incubated 25 min at 21°C to allow the worms to settle. Then, basal activity was measured for 30 minutes to normalize the movement activity in each well at the beginning of the assay. Immediately after drug treatment, each score activity was recorded for a 120-minute period. Motility was calculated according to the formula:

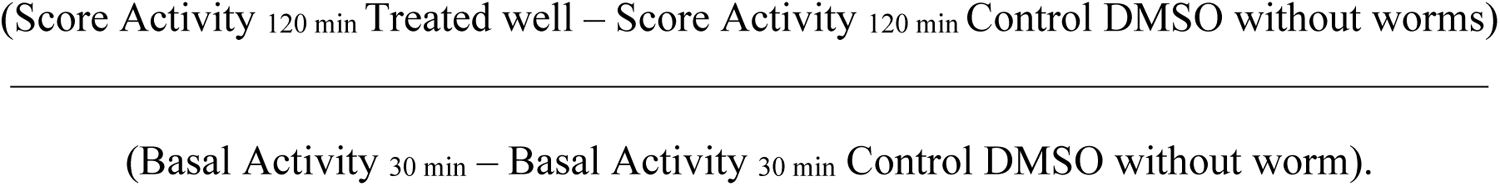

Motility percentages were calculated for each treated well as –fold induction relative to DMSO treated worms which was set to 100.

#### 2.4.2 LMA on the *H. contortus* isolates

The LMA on *H. contortus* measures the potency of anthelmintics in inhibiting motility in infective L3 larvae (iL3). Prior to experiment, *H. contortus* larvae were treated to discard cuticle. Briefly, worms were incubated 20 minutes at 37°C, in tap water supplemented by NaCl 0.15 % and vigorously shacked by vortex every 5 minutes. To prevent larval aggregation, larvae were filtered through a 40 µm mesh in LB medium. Each well of a 96-flat well plate received 80 iL3 suspended in a 200 µl final volume of LB medium. Then, the plates were treated with compounds and subsequently incubated at 37°C for 24 hours within a humidified incubator, maintaining a 5% CO_2_ atmosphere and humidity levels ≥ 90%. Following the incubation period, motility of the larvae (iL3) was restored by exposing them to light at room temperature for 5 minutes. Thereafter, the movement of worms within each well was recorded over a 15-minute duration using WMi technology. The motility of worms in each well was then standardized against the average motility of control wells to derive the motility inhibition values (%). Post log_10_-transformation of compound concentrations, dose-response curves for the motility assay were fitted using a sigmoidal model with variable slope parameters. The 95% confidence limits were determined and graphed using GraphPad Prism 6 software package. IC50 values, i.e., the concentration at which 50% of the worms are immobilized by the drug were then calculated (GraphPad, San Diego, CA, USA). Each value was the mean of triplicate and the experiments were performed at least 3 times.

### 2.5 Statistical analysis

Firstly, the bioassay data were tested for normality using the D’Agostino & Pearson omnibus normality test. Thereafter, a two-way analysis of variance (ANOVA) was used to compare the effect of two categorical variables on the motility of worms: (i) biological condition (referring to drug tolerance status of strains; control, resistant or hypersensitive) and (ii) drug treatment condition (IVM, MOX and EPR) at different concentrations. Tuckey’s test was applied for multiple comparisons. Biological repeats are defined as independent experiments conducted on separate populations of animals on different days. These analyses were performed using GraphPad Prism6 software package.

## 3. Results

### 3.1 Motility assay to assess IVM efficacy in adult *C. elegans*

Different conditions were initially tested to obtain an optimal *C. elegans* motility response. We first verified that DMSO, used as a compound vehicle did not affect adult *C. elegans* motility when used at concentrations below 0.5%. Subsequently all experiments were conducted in medium with DMSO under 0.5%. We then determined the number of worms required for the experiment and established a linear correlation with motility between 30 and 90 worms (data not shown). The experiments were then performed with 40 worms per well, in the linearity range, and the motility was recorded during 120 minutes. We then used worm motility test to compare IVM susceptibility of several *C. elegans* strains: the wild-type Bristol N2 (N2B), IVM-selected (IVR10) and *nhr-8* loss-of-function (AE501; *nhr8(ok186)*) (Fig. 1). Fig. 1A shows representative images of worms after 120 minutes incubation without or with increasing concentration of IVM. Images clearly show nice curved animals in control (DMSO) or at 0.01 µM of IVM. In these conditions, animals were regularly moving revealing that such a low concentration of IVM did not affect worm motility whatever the *C. elegans* strain. IVM at 1 µM was able to induce total population body stiffness phenotype representing complete altered worm motility in all strains. Images reveal clear visible differences in motility phenotype between strains at 0.1 µM of IVM. All N2B and *nhr-8* deficient worms being immobile, while a significant proportion of IVM-resistant worms were still moving. To quantify such differences, dose-response curves for IVM toward motility of young adult *C. elegans* of the three strains were graphed and are presented in Fig. 1B. IC50s, i.e., the concentration of IVM at which 50% of animals are immobile and resistance factor (RF) values, reflecting differences in the IC50 compared with that of N2B, are shown in Table 1.

**Fig 1.**
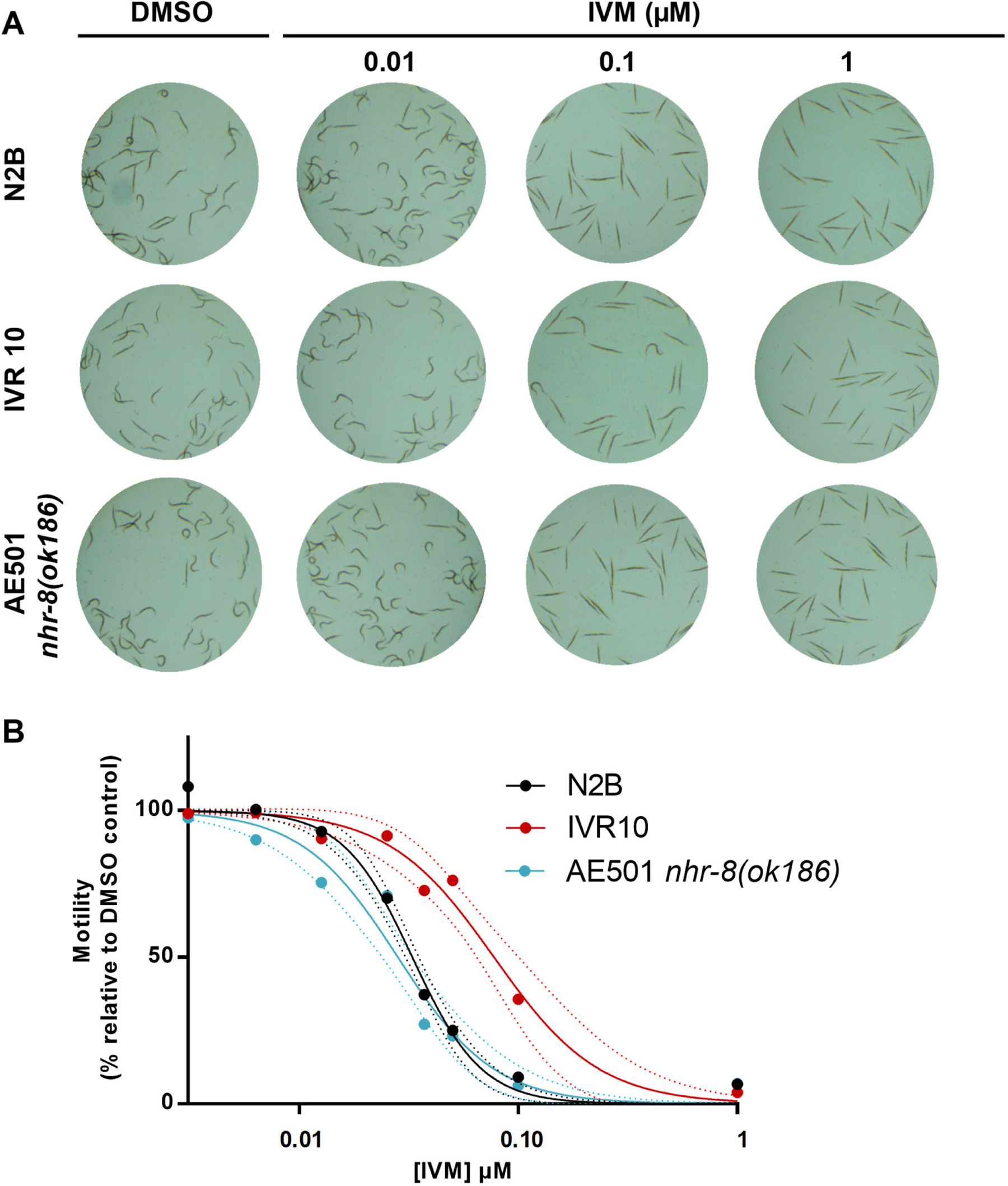
Comparison of IVM efficacy in *C. elegans* strains using larval motility assay. **(A)** Top view of *C. elegans* adults in representative wells of 96-well plates after 120 minutes exposure without or with IVM at 0.01, 0.10 and 1.00 µM. **(B)** Representative concentration-response curve of *C. elegans* motility inhibition after exposure to IVM. Worm motility was assessed using the wMicroTracker which allows to quantify the locomotor activity of a worm population. Young adults of wild-type Bristol N2 (N2B, black), IVM resistant (IVR10, red) and IVM hypersensitive (AE501 *nhr(ok186),* blue) strains were incubated for 120 minutes at 21°C in the presence of increasing concentrations of IVM (0.0125 to 1 µM). For each strain, motility percentages are expressed as –fold induction relative to DMSO treated worms, which is set to 100, and are reported as the mean and 95% confidence bands (dotted lines), a triplicates per conditions out of 5-20 experiments.

**TABLE 1.**
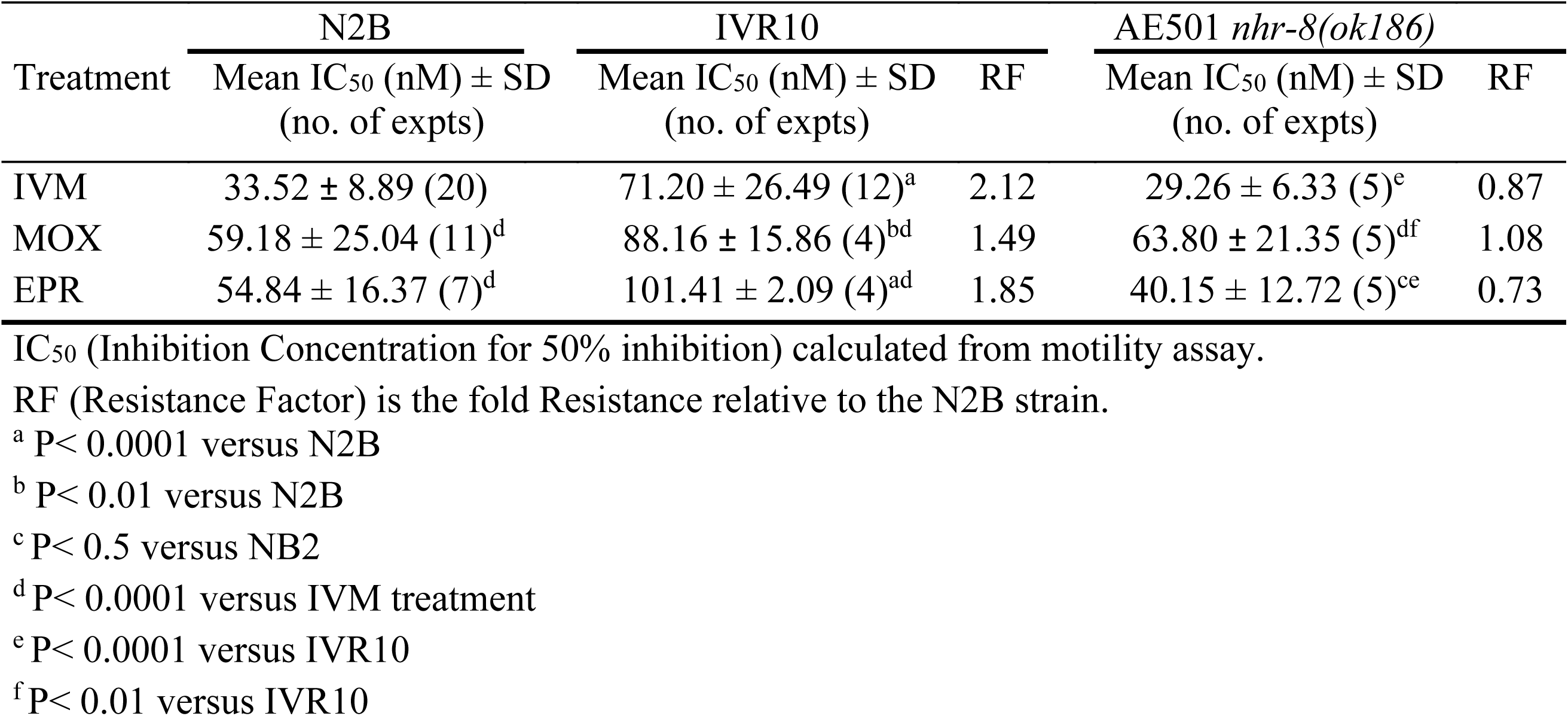
Effect of IVM, MOX and EPR on worm motility in wild-type N2B, IVM-selected (IVR10) and IVM-hypersensitive *nhr-8*-deficient (AE501 *nhr-8(ok186)*) *C. elegans* strains.

In agreement with the images, dose-response curves showed that IVM was able to alter worm motility of all strains. However, IVM displayed different potencies in affecting *C. elegans* motility. Indeed, similar potency of IVM to inhibit motility was observed for N2B and *nhr-8* deficient worms as shown by the superposition of the two dose-response curves and by the comparable EC50 values (33.52 ± 8.89 nM and 29.26 ± 6.33 nM, respectively). In contrast, the dose-response curve was significantly shifted to the right for the IVM-selected strain, reflecting a decrease in susceptibility to IVM. Indeed, IC50 of IVM was 2.12-fold higher for IVR10 than wild-type (71.20 ± 26.49 nM and 33.52 ± 8.89 nM respectively, P<0.0001) in agreement with a higher tolerance of IVR10 worms to IVM when compared with wild-type N2B (Ménez et al., 2016)

### 3.2 Impact of MOX and EPR on adult *C. elegans* motility

Because cross-resistance between MLs is well described, we then investigated the ability of the motility test to discriminate ML tolerance between strains, by studying MOX and EPR impact on the motility of each strain. Table 1 shows that IVR10 strain was more tolerant to the three drugs tested compared with the wild-type strain with IVM being the most potent drug. Indeed, IVR10 strain was 2.12-fold, 1.85-fold and 1.49-fold less sensitive to IVM, EPR and MOX respectively when compared with N2B strain. IVM was also the most potent drug for N2B strain (Table 1). The trend towards greater IVM efficacy on worm motility phenotype seems to be continuing also in *nhr-8* deficient worm, however differences are not significant compared with EPR treatment. The less potent drug toward motility in this strain was MOX. Taken together, these results clearly show that the motility assay is suitable to evaluate ML toxicity and to discriminate drug resistant from susceptible *C. elegans* strains but not appropriate to discriminate NHR-8 deficient worms, previously described as hypersensitive to IVM (Ménez et al., 2019).

### 3.3 Motility assay to discriminate susceptible from resistant *H. contortus* in the field

To respond to the urgent need of veterinarians and farmers to confirm resistance, in the context of treatment failure and suspicion of drug resistance in the field, we decided to adapt the use of WMicrotracker^TM^ based on previous work (Taki et al., 2021), to monitor motility in *H. contortus* and discriminate susceptible from resistant isolates. The results of the dose-response experiments with IVM, MOX and EPR against both susceptible and resistant *H. contortus* isolates are shown in Fig 2.

**Fig 2.**
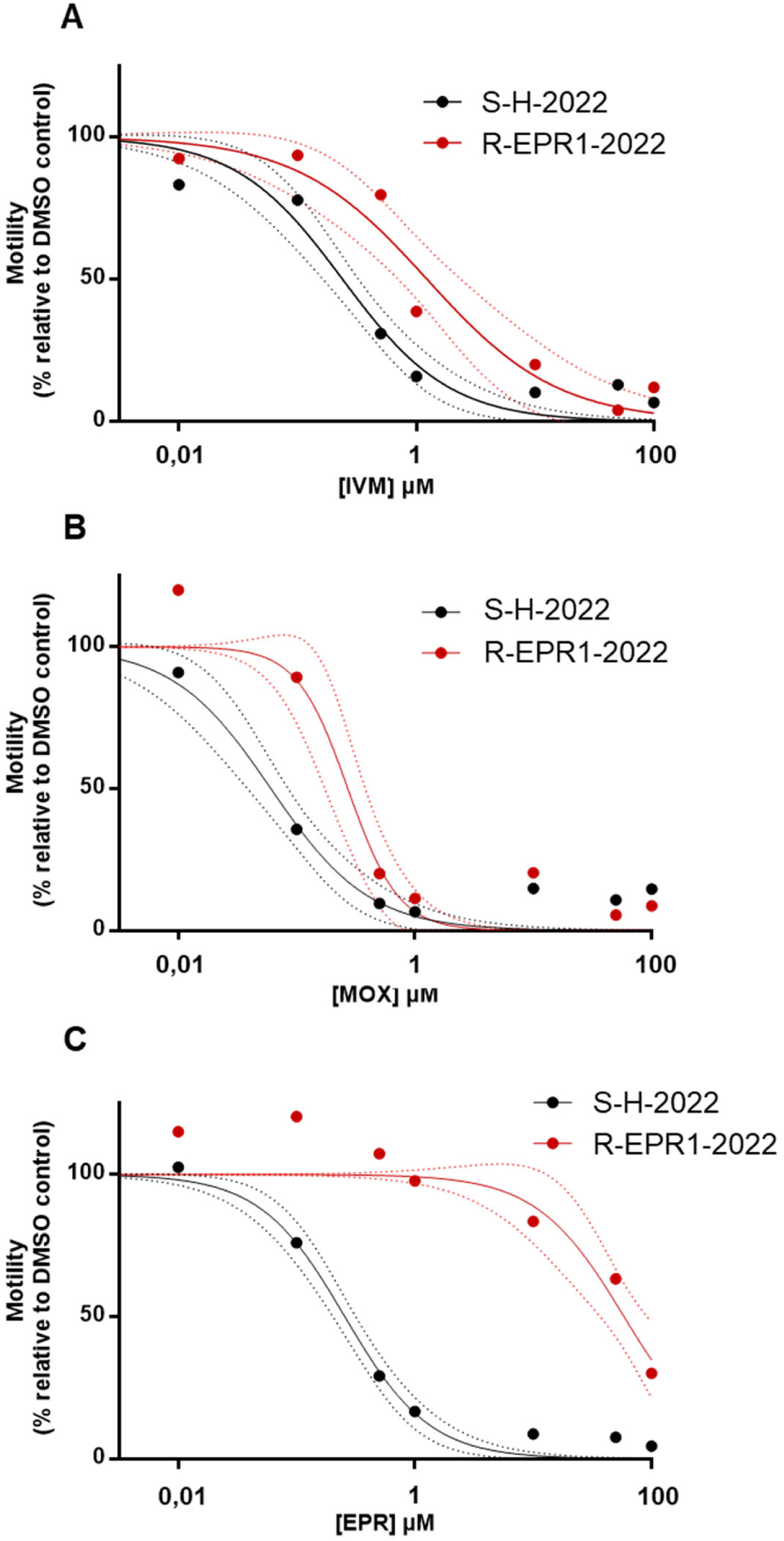
Comparison of ML susceptibility of the LM-susceptible (S-H-2022) and LM-resistant (R-EPR1-2022) *H. contortus* strains in a larval motility assay. Representative concentration-response curve of *H. contortus* motility inhibition following exposure to **(A)** IVM, **(B)** MOX and **(C)** EPR. Worm motility was assessed using the WMicroTracker which allows to quantify the locomotor activity of a worm population. IL3 of susceptible (S-H-2022, black), LMs resistant (R-EPR1-2022, red) strains were incubated for 120 minutes at 21°C in the presence of increasing concentrations of IVM (0.01 to 100 µM). For each strain, motility percentages are expressed as –fold induction relative to DMSO, which is set to 100, and are reported as the mean and 95% confidence bands (dotted lines), a triplicates per conditions out of 3-7experiments

**Figure 3:**
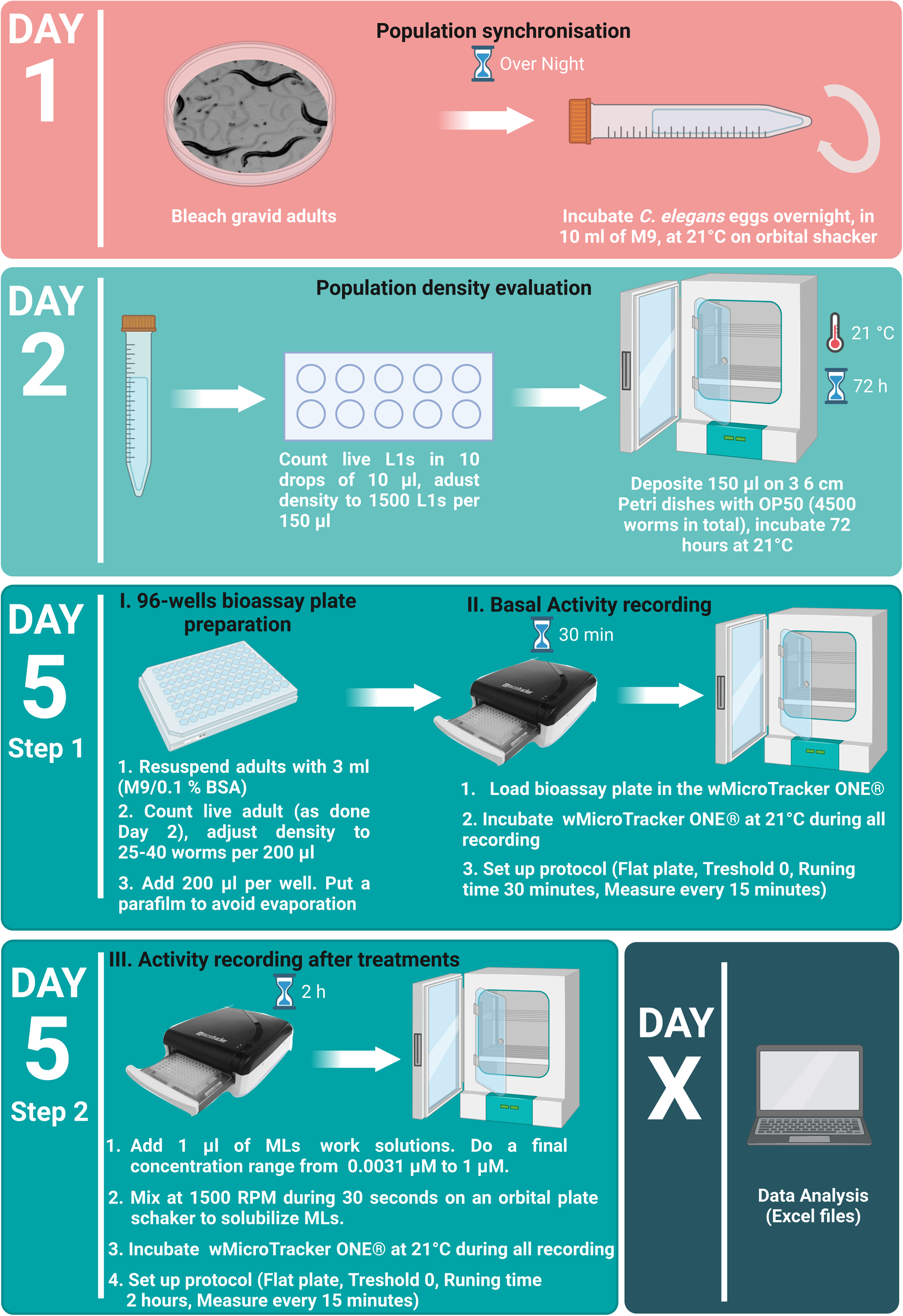
Caenorhabditis elegans experimental setup. Schematic overview of the experimental procedure and associated timeline for culturing C. elegans. The timeline for preaparing bioassay plate for larval motility measurement using WomMicrotracker is also provided. To prevent adult sticking, all plastic material in contact with adult larvae (Step 5) must be impregnated with 0.1% BSA (Bovine Serum Albumin). **B**SA: Bovine Serum Albumine). The figure is inspied by Figure from Preez et al, 2020. Created with BioRender.com.

**Figure 4:**
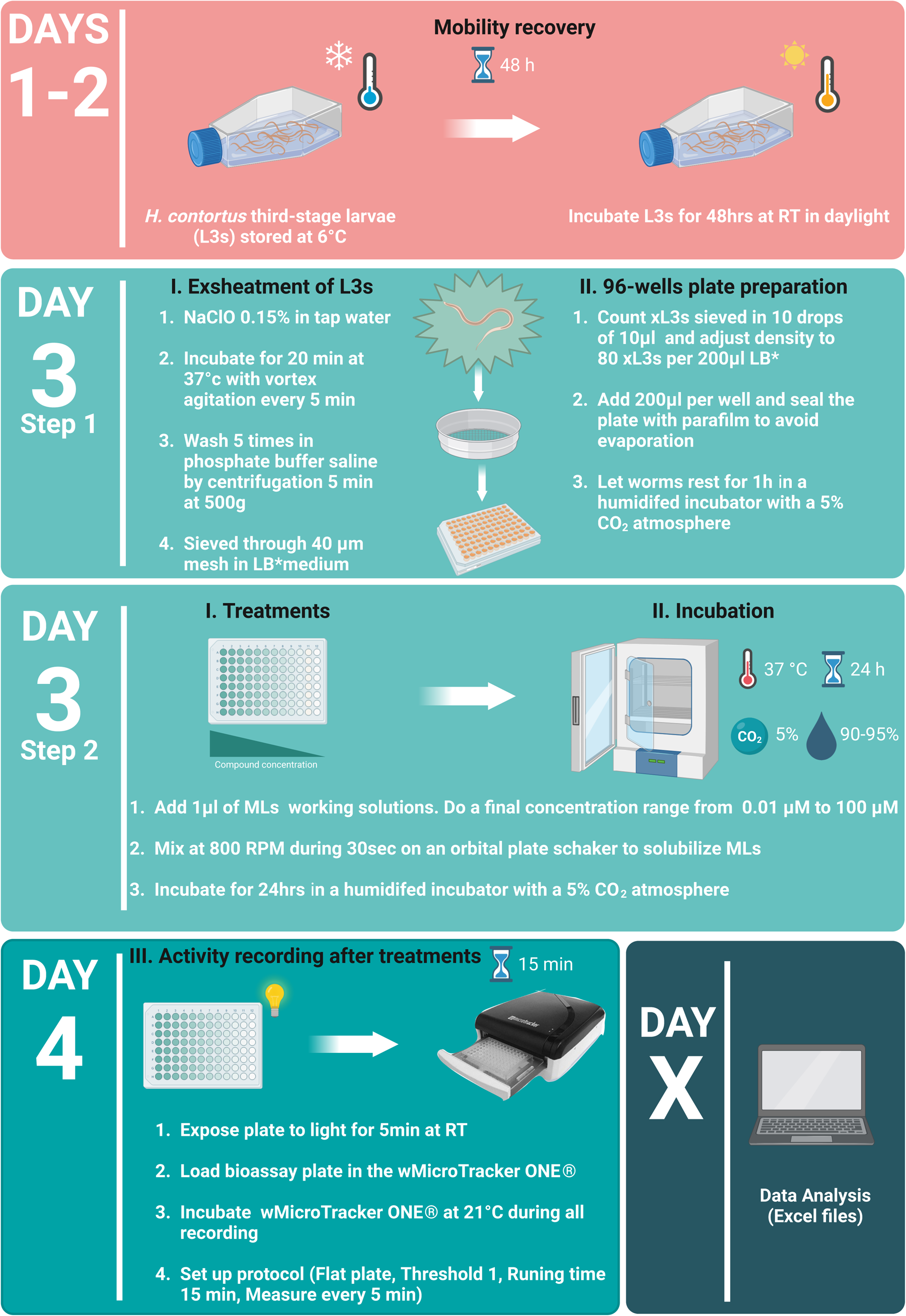
Haemonchus contortus experimental setup. Schematic overview of the experimental procedure and associated timeline for or larval motility measurement using WomMicrotracker. To prevent sticking, all plastic material in contact with the larvae must be impregnated with 0.1% BSA (Bovine Serum Albumin). LB* : Luria Bertani medium supplemented with final concentrations of 100IU/mL of penicillin, 100μg/mL of streptomycin and 0.25μg/mL of amphotericin B. The figure is inspied by Figure from Preez et al, 2020. Created with BioRender.com.

As expected, increasing concentrations of all drugs were able to alter worm motility of both strains. However, there was a significant difference on their potency. IC50 values for each drug on both isolates have been calculated and are presented in Table 2.

**TABLE 2.** Effect of IVM, MOX and EPR on worm motility in MLs-susceptible (S-H-2022) and MLs-resistant (R-EPR1-2022) *H. contortus* isolates.

|  | S-H-2022 | R-EPR1-2022 |  |
| --- | --- | --- | --- |
| Treatment | Mean IC<sub>50</sub> (μM) ± SD<br>(no. of expts) | Mean IC<sub>50</sub> (μM) ± SD<br>(no. of expts) | RF |
| IVM | 0.21 ± 0.12 (7) | 1.10 ± 0.42 (3)<sup>a</sup> | 5.24 |
| MOX | 0.05 ± 0.04 (6) | 0.27 ± 0.06 (3)<sup>bc</sup> | 5.40 |
| EPR | 0.26 ± 0.09 (7) | 60.89 ± 16.92 (4)<sup>ad</sup> | 234.19 |

IC<sub>50</sub> (Inhibition Concentration for 50% inhibition) calculated from motility assay.
RF (Resistance Factor) is the fold Resistance relative to S-H-2022.
<sup>a</sup> P< 0.0001 versus S-H-2022
<sup>b</sup> P< 0.001 versus S-H-2022
<sup>c</sup> P< 0.05 versus IVM treatment
<sup>d</sup> P< 0.0001 versus IVM treatment

Firstly, all three drugs were more effective on the susceptible strain than on the resistant strain, as demonstrated by the lower IC_50_ values. Secondly, on both isolates, MOX was the most potent drug when compared to IVM and EPR as shown by the smaller IC_50_ within isolates (, P<0.0001 versus IVM treatment.). However, the degree of resistance to MOX for the resistant isolate was identical to that of IVM as demonstrated by the same resistance factor (RF of 5.24 and 5.40 for IVM and MOX, respectively, P<0.0001 versus susceptible isolate). The most substantial degree of resistance of the resistant isolate was observed for EPR. Indeed, the RF for this molecule was 234 and is reflected by a huge shift of the curve to the right. Indeed, EPR was the least potent compared with the two other drugs, while IVM displayed intermediate potency.

## 4. Discussion

To establish robust nematode control programs, it is essential to integrate highly sensitive and easy methods for detecting and regularly monitoring anthelmintic resistance. However, in practical farm applications, the available assays are often laborious. Reduced motility is a key phenotype for evaluating the bioactivity of anthelmintic compounds. In this study, we sought to evaluate the feasibility of measuring the worm motility assay using the automated apparatus, WMicrotracker^TM^, as a reliable, rapid, and high-throughput method for assessing ML susceptibility in nematodes. Our final objective was to discriminate between susceptible and resistant nematodes. We demonstrated, for the first time, the reproducibility of this method, showcasing its sensitivity in differentiating between susceptible and resistant isolates of *C. elegans* and *H. contortus*.

The primary objective of our investigation was to conduct a comparative analysis of the efficacy of the anthelmintic macrocyclic lactones IVM, MOX and EPR on both susceptible and resistant strains of the nematode model *C. elegans*, as well as the field parasite *H. contortus*.

We first explored the efficacy of the motility assay to distinguish between susceptible and resistant strains of *C. elegans*. Traditionally, the larval development assay (LDA) was the reference test for assessing ML efficacy in *C. elegans.* Subsequently, we explored the potential application of the worm motility assay within the nematode model *C. elegans*. In this study, we conducted comparative analyses of the impact of MLs on worm motility inhibition across three strains: the wild-type Bristol N2 strain (N2B), the IVM-selected strain IVR10 and the IVM-hypersensitive *nhr-8* deficient strain. While the WMi assay has been previously employed to monitor ML efficacy in *C. elegans*, this study marks the first instance where its utility was explored to compare drug resistant and susceptible strains. As expected, IVM, MOX and EPR were able to alter worm motility of all strains studied and differences in potency of the drugs were observed between strains. Indeed, IVM was able to alter N2B worm motility with very high potency, while MOX and EPR showed comparable potency, lower than IVM. We then measured the impact of ML on worm motility of ML-resistant *C. elegans*. This strain has been selected on IVM pressure and is highly resistant to the three MLs, as determined by LDA (Ménez et al., 2016). Significantly higher concentrations of the three ML compounds were required to affect the worm motility of the IVM-resistant strains, showing that IVR10 strain was not only tolerant to IVM but also to MOX and to EPR, as consistently observed in the LDA data. Interestingly, our results indicated equal potency of IVM in inhibiting the motility of both the parental N2B strain and the IVM-hypersensitive strain, as evidenced by their identical resistance factors (RFs). However, while by using LDA, *nhr-8* deficient strain shows hyper susceptibility to IVM (Ménez et al., 2019), WMi test did not reveal a differential susceptibility to IVM of the *nhr-8* deficient strain when compared to N2B, indicating a potential limitation of the motility test. Nonetheless, the WMi assay proved effective in discriminating between N2B and IVM-resistant strains. Comparative analyses of the efficacy of the three drugs revealed IVM as the most potent drug in the N2B and IVR10 strains using WMi. This aligns with previous findings demonstrating the sensitivity of N2 Bristol motility to IVM and MOX (Ardelli et al., 2009). However, while the trend towards greater IVM efficacy persisted in *nhr-8* deficient worm, differences were not significant compared with EPR treatment (Table 2). This reveals differences with previous LDA data, showing a higher efficacy of EPR in IVR10 worms compared with N2B. This is certainly due to the drug effect on the phenotype observed which is different from one test to another. Motility is affected by ML in a different way than larval development. This reveals discrepancies between tests and encourage one to take into account the assays accuracy when choosing a test to evaluate drug efficacy. MLs, and especially IVM, are well characterized for its inhibitory effect on worm motility. Nevertheless, disparities in concentration with other studies were observed. Table 3 summarizes IC50s obtained with different phenotypic assays in *C. elegans*. Indeed, our results showed a significant motility inhibition at ML concentrations lower than those used in previous studies. As an example, Hahnel *et al*. conducted a comparative analysis of the potency of IVM and MOX in suppressing N2B motility by using WMi. Their findings revealed that MOX exhibited 2.4 times greater potency in inhibiting motility compared to IVM in their experiment (Hahnel et al., 2021). In the same range, Risi *et al* tested the effect of IVM on *C. elegans* L4 motility and obtained an IC50 value of 190 nM (Risi et al., 2019). Such discrepancies could be attributed to the specific method employed for the application of ML on the worm, which account for drug bioavailability in the assay. In our approach, we directly introduce 1 µl of ML into the well without prior dilution in another container. The reduced bioavailability of ML in some studies may be associated with the retention of ML in plastic, and bioavailability is contingent on this factor. We recommend avoiding ML contact with multiple containers to improve ML availability during assay. Table 3 further highlights that the IC50 values are notably lower for LDA than those obtained from motility tests. It could be explained, as illustrated by Munguia *et al*, that the extended read-out times of the LDAs in *H. contortus* resulted in notably heightened sensitivity to standard anthelmintics, including IVM, compared to a 24-hour test directly measuring the effects of anthelmintics on L3 motility (Munguía et al., 2022).

**TABLE 3.** Comparison of Macrocyclic lactones IC50s (nM) obtained by different *C. elegans* phenotypic assay.

|  |  | C. elegans strains |  |  |  |  |  |  |  |  |  |  |
| --- | --- | --- | --- | --- | --- | --- | --- | --- | --- | --- | --- | --- |
| Assay | Informations about experiments | N2B |  |  | NHR-8 |  |  |  | IVR10 |  |  |  |
|  |  | IVM | MOX | EPR | IVM | MOX | EPR |  | IVM | MOX | EPR | References |
| Motility assay | Adults, Worm Microtracker | 33,52 | 59,18 | 54,84 | 29,26 | 63,80 | 40,15 |  | 71,20 | 88,16 | 101,41 | Current study |
|  | L4, Worm Microtracker | 190 | ND | ND | ND | ND | ND |  | ND | ND | ND | (Risi et al., 2019) |
|  | Adults, Worm Microtracker | 290 | 120 | ND | ND | ND | ND |  | ND | ND | ND | (Hahnel et al., 2021) |
|  | L4, Worm Microtracker | 150-500 | ND | 100-300 | ND | ND | ND | ND | ND | ND | ND | (Suárez et al., 2022) |
| Larval development assay | Agar plates | 1,69 | 1,77 | 1,19 | ND | ND | ND |  | 12,43 | 3,06 | 17,82 | (Ménez et al., 2016) |
|  | Agar plates | 1,63 | ND | ND | 0,96 | ND | ND |  | ND | ND | ND | (Ménez et al., 2019) |
| Pharynx pumping | 8-channel chip | 1420 | 900 | ND | ND | ND | ND |  | ND | ND | ND | (Hahnel et al., 2021) |
|  | ScreenChip | 51 | 42 | ND | ND | ND | ND |  | ND | ND | ND | (Hahnel et al., 2021) |

Mean IC<sub>50</sub> (Inhibition Concentration for 50% inhibition) calculated by authors (See References)
ND (Not Determined)

Taken together, these results definitively validate the efficacy of the motility assay using WMi in assessing ML effectiveness in *C. elegans*. Notably, this study marks the inaugural instance wherein the results demonstrate the capability of WMi to distinguish between drug-resistant and susceptible *C. elegans* strains. However, while the LDA effectively discerns hypersensitivity in *nhr-8* deficient worms, the WMi assay did not reveal such heightened sensitivity to the drug. Nevertheless, our study highlights the complexity of interpreting ML concentration responses across various studies, owing to variations in drug application methods, phenotypes assessed, and other contributing factors (time of incubation, *C. elegans* development stage…).

While the fecal egg count reduction test (FECRT) is presently the favored approach for identifying anthelmintic resistance at the farm level, it bears the limitations of being labor-intensive, costly, and capable of detecting resistance only when it has already reached relatively high levels. *In vitro* assays are regarded as the most efficient for the early detection of anthelmintic resistance. In this context, monitoring worm motility via the larval migration assay in *H. contortus* has been suggested (Demeler et al., 2012; Dolinská et al., 2016). However, the manual counting of larvae, which is labor-intensive, could be a reason why this test isn’t commonly employed in the field for detecting drug resistance.

Therefore, we studied WMi application on *H. contortus* L3 larvae and compared ML efficacy in two isolates, one susceptible and one, collected in farm, suspected of being ML resistant. Indeed, the IC_50_ values obtained for IVM and MOX (Table 2) were consistent with the concentrations previously reported for motility assays in xL3s after 24 hours of incubation (Munguía et al., 2022; Taki et al., 2021). Then, lower efficacy was observed for the three drugs in the resistant isolates. The resistance factor, for the 3 MLs, between susceptible and resistant isolates of *H. contortus* was consistently greater than 5, indicating a significant level of resistance to these 3 drugs. MOX was more potent than IVM, while EPR was the least potent drug, displaying the highest resistance factor compared to IVM and MOX (RF up to 234). However, IVM and MOX displayed the same RF (5.24 and 5.40 respectively). The observed order of potency of MLs for *H. contortus* R-EPR1-2022 closely mirrored that observed in the resistant *H. contortus* Kokstad isolate, highlighting MOX as significantly more potent than IVM and EPR (Kotze et al., 2006; Ménez et al., 2016). To our knowledge, this is the first report showing the significance of high-throughput quantitative motility assessment using WMi in detecting anthelmintic resistance in a field parasite. Professionals want an easy, fast and high throughput test applicable on the field. Interestingly, in just a four-day period on the L3 stage and with minimum human intervention, our results showed first that we were able to evaluate and compare ML efficacy on susceptible *H. contortus* isolates. Furthermore, our groundbreaking findings reveal that the worm motility assay, using microtracker technology, could offer a rapid, efficient, and highly practical method for assisting veterinarians and farmers in the field in detecting anthelmintic resistance on *H. contortus* isolates. Few studies have explored the motility test as an effective method for detecting resistance to anthelmintic. As an example, a report suggested that motility of the L3 stage was a poor phenotype for detecting and measuring anthelmintic resistance of different gastrointestinal nematodes (George et al., 2018). Indeed, in this study, resistance factor never achieved more than 2 in score whatever the parasite being tested. These differences may be explained by (i) the test used which was based on larval migration less sensitive than the motility assay used in the present study and (ii) the presence of the cuticle as authors worked on sheathed L3 which is a robust barrier, shielding the worm from its environment, especially against xenobiotics (Page et al., 2014). We conducted our research on exsheathed L3 larvae, which guarantee that both susceptible and resistant L3s were optimally exposed to the drug. Another work on the filarial nematode *Dirofilaria immitis*, has shown that motility of microfilaria was not a reliable phenotype for detecting resistance in this parasite, encouraging professional to *in vivo* assay such as the microfilaria suppression test in the presence of a suspect case of resistant isolates (Maclean et al., 2017). Additional investigations are needed to explore its potential broader applications to other parasites.

In conclusion, our findings highlight for the first time, that the WMicrotracker motility assay stands as a robust test to be employed for effectively discerning between ML-susceptible and resistant nematodes. This distinction is achieved within an efficient high-throughput timeframe of just 4 to 5 days. Moreover, this test could be a valuable tool to detect drug resistance in *H. contortus* as it allows to (i) discriminate ML susceptible from resistant isolates of *H. contortus* from the field, (ii) show cross resistance to the three molecules in both susceptible and resistant isolates and (iii) highlight a huge resistance to EPR consistent with the failure of treatment reported from the field.

## Acknowledgments

This work was funded by the France Futur Elevage (F2E Institut-Carnot Santé Animale) ‘Antherin’ project. Figure 3 and 4 were created with BioRender.com. We sincerely thank Professor Roger K. Prichard for engaging in fruitful discussions, conducting a meticulous review, and providing invaluable feedback on the manuscript.

## Author Contributions

The research was conceptualized and the experiments were designed by M. A. and A. L. Funding acquisition was performed by M. A. and A. L. Most of the experiments were conducted by M. A. and M. G. with punctual contributions of J. P. and C. B. The animal experiments to provide *H. contortus* isolates was performed by J. P., S. J. and P. J. M. A. conducted the data analysis, the statistical analysis and drafted the manuscript, with specific contributions and input from all authors.

## Additional Information

Competing Interests: The authors declare no competing interests.

## Notes

### Competing Interest Statement

The authors have declared no competing interest.

## References

Ardelli, B.F., Stitt, L.E., Tompkins, J.B., Prichard, R.K., 2009. A comparison of the effects of ivermectin and moxidectin on the nematode Caenorhabditis elegans. Vet. Parasitol. 165, 96–108. 10.1016/j.vetpar.2009.06.043

Bianchi, J.I., Stockert, J.C., Buzz, L.I., Blázquez-Castro, A., Hernán Simonetta, S., 2015. Reliable screening of dye phototoxicity by using a Caenorhabditis elegans fast bioassay. PLoS One 10, 1–15. 10.1371/journal.pone.0128898

Bourguinat, C., Lee, A.C.Y., Lizundia, R., Blagburn, B.L., Liotta, J.L., Kraus, M.S., Keller, K., Epe, C., Letourneau, L., Kleinman, C.L., Paterson, T., Carretón, E., Montoya-Alonso, J.A., Smith, H., Bhan, A., Peregrine, A.S., Carmichael, J., Drake, J., Schenker, R., Kaminsky, R., Bowman, D.D., Geary, T.G., Prichard, R.K., 2015. Macrocyclic lactone resistance in Dirofilaria immitis: Failure of heartworm preventives and investigation of genetic markers for resistance. Vet. Parasitol. 210, 167–178. 10.1016/j.vetpar.2015.04.002

Buckingham, S.D., Partridge, F.A., Sattelle, D.B., 2014. Automated, high-throughput, motility analysis in Caenorhabditis elegans and parasitic nematodes: Applications in the search for new anthelmintics. Int. J. Parasitol. Drugs Drug Resist. 4, 226–232. 10.1016/j.ijpddr.2014.10.004

Demeler, J., Kleinschmidt, N., Küttler, U., Koopmann, R., von Samson-Himmelstjerna, G., 2012. Evaluation of the Egg Hatch Assay and the Larval Migration Inhibition Assay to detect anthelmintic resistance in cattle parasitic nematodes on farms. Parasitol. Int. 61, 614–618. 10.1016/j.parint.2012.06.003

Dolinská, M.U., Königová, A., Babják, M., Várady, M., 2016. Comparison of two in vitro methods for the detection of ivermectin resistance in Haemonchus contortus in sheep. Helminthol. 53, 120–125. 10.1515/helmin-2015-0002

Doyle, S.R., Bourguinat, C., Nana-Djeunga, H.C., Kengne-Ouafo, J.A., Pion, S.D.S., Bopda, J., Kamgno, J., Wanji, S., Che, H., Kuesel, A.C., Walker, M., Basáñez, M.G., Boakye, D.A., Osei-Atweneboana, M.Y., Boussinesq, M., Prichard, R.K., Grant, W.N., 2017. Genome-wide analysis of ivermectin response by Onchocerca volvulus reveals that genetic drift and soft selective sweeps contribute to loss of drug sensitivity, PLoS Neglected Tropical Diseases. 10.1371/journal.pntd.0005816

Driscoll, M., Dean, E., Reilly, E., Bergholz, E., Chalfie, M., 1989. Genetic and molecular analysis of a Caenorhabditis elegans beta-tubulin that conveys benzimidazole sensitivity. J. Cell Biol. 109, 2993–3003. 10.1083/jcb.109.6.2993

European Medicines Agency, 2016. Reflection paper on anthelmintic resistance (Draft 2). Eur. Med. Agency 44, 1–16.

Fitzpatrick, J.L., 2013. Global food security: The impact of veterinary parasites and parasitologists. Vet. Parasitol. 195, 233–248. 10.1016/j.vetpar.2013.04.005

Garbin, V.P., Munguía, B., Saldaña, J.C., Deschamps, C., Cipriano, R.R., Molento, M.B., 2021. Chemical characterization and in vitro anthelmintic activity of Citrus bergamia Risso and Citrus X paradisii Macfad essential oil against Haemonchus contortus Kirby isolate. Acta Trop. 217. 10.1016/j.actatropica.2021.105869

George, M.M., Lopez-Soberal, L., Storey, B.E., Howell, S.B., Kaplan, R.M., 2018. Motility in the L3 stage is a poor phenotype for detecting and measuring resistance to avermectin/milbemycin drugs in gastrointestinal nematodes of livestock. Int. J. Parasitol. Drugs Drug Resist. 8, 22–30. 10.1016/j.ijpddr.2017.12.002

Hahnel, S.R., Dilks, C.M., Heisler, I., Andersen, E.C., Kulke, D., 2020. Caenorhabditis elegans in anthelmintic research – Old model, new perspectives. Int. J. Parasitol. Drugs Drug Resist. 14, 237–248. 10.1016/j.ijpddr.2020.09.005

Hahnel, S.R., Roberts, W.M., Heisler, I., Kulke, D., Weeks, J.C., 2021. Comparison of electrophysiological and motility assays to study anthelmintic effects in Caenorhabditis elegans. Int. J. Parasitol. Drugs Drug Resist. 16, 174–187. 10.1016/j.ijpddr.2021.05.005

James, C.E., Davey, M.W., 2009. Increased expression of ABC transport proteins is associated with ivermectin resistance in the model nematode Caenorhabditis elegans. Int. J. Parasitol. 39, 213–220. 10.1016/j.ijpara.2008.06.009

Jouffroy, S., Bordes, L., Grisez, C., Sutra, J.F., Cazajous, T., Lafon, J., Dumont, N., Chastel, M., Vial-Novella, C., Achard, D., Karembe, H., Devaux, M., Abbadie, M., Delmas, C., Lespine, A., Jacquiet, P., 2023. First report of eprinomectin-resistant isolates of Haemonchus contortus in 5 dairy sheep farms from the Pyrénées Atlantiques département in France. Parasitology 150, 365–373. 10.1017/S0031182023000069

Kotze, A.C., Le Jambre, L.F., O’Grady, J., 2006. A modified larval migration assay for detection of resistance to macrocyclic lactones in Haemonchus contortus, and drug screening with Trichostrongylidae parasites. Vet. Parasitol. 137, 294–305. 10.1016/j.vetpar.2006.01.017

Laing, R., Gillan, V., Devaney, E., 2017. Ivermectin – Old Drug, New Tricks? Trends Parasitol. 33, 463–472. 10.1016/j.pt.2017.02.004

Liu, M., Kipanga, P., Mai, A.H., Dhondt, I., Braeckman, B.P., De Borggraeve, W., Luyten, W., 2018. Bioassay-guided isolation of three anthelmintic compounds from Warburgia ugandensis Sprague subspecies ugandensis, and the mechanism of action of polygodial. Int. J. Parasitol. 48, 833–844. 10.1016/j.ijpara.2017.11.009

Liu, M., Landuyt, B., Klaassen, H., Geldhof, P., Luyten, W., 2019. Screening of a drug repurposing library with a nematode motility assay identifies promising anthelmintic hits against Cooperia oncophora and other ruminant parasites. Vet. Parasitol. 265, 15–18. 10.1016/j.vetpar.2018.11.014

Liu, M., Lu, J.G., Yang, M.R., Jiang, Z.H., Wan, X., Luyten, W., 2022. Bioassay-Guided Isolation of Anthelmintic Components from Semen pharbitidis, and the Mechanism of Action of Pharbitin. Int. J. Mol. Sci. 23. 10.3390/ijms232415739

Maclean, M.J., Savadelis, M.D., Coates, R., Dzimianski, M.T., Jones, C., Benbow, C., Storey, B.E., Kaplan, R.M., Moorhead, A.R., Wolstenholme, A.J., 2017. Does evaluation of in vitro microfilarial motility reflect the resistance status of Dirofilaria immitis isolates to macrocyclic lactones? Parasites and Vectors 10, 17–23. 10.1186/s13071-017-2436-6

Martin, R.J., Robertson, A.P., Choudhary, S., 2020. Trends in Ivermectin : An Anthelmintic, an Insecticide, and Much More IvermecƟn analogs. Trends Parasitol. 1–17. 10.1016/j.pt.2020.10.005

Ménez, C., Alberich, M., Courtot, E., Guegnard, F., Blanchard, A., Aguilaniu, H., Lespine, A., 2019. The transcription factor NHR-8: A new target to increase ivermectin efficacy in nematodes, PLOS Pathogens. 10.1371/journal.ppat.1007598

Ménez, C., Alberich, M., Kansoh, D., Blanchard, A., Lespine, A., 2016. Acquired tolerance to ivermectin and moxidectin after drug selection pressure in the nematode Caenorhabditis elegans. Antimicrob. Agents Chemother. 60, 4809–4819. 10.1128/AAC.00713-16

Morgan, E.R., Aziz, N.A.A., Blanchard, A., Charlier, J., Charvet, C., Claerebout, E., Geldhof, P., Greer, A.W., Hertzberg, H., Hodgkinson, J., Höglund, J., Hoste, H., Kaplan, R.M., Martínez-Valladares, M., Mitchell, S., Ploeger, H.W., Rinaldi, L., von Samson-Himmelstjerna, G., Sotiraki, S., Schnyder, M., Skuce, P., Bartley, D., Kenyon, F., Thamsborg, S.M., Vineer, H.R., de Waal, T., Williams, A.R., van Wyk, J.A., Vercruysse, J., 2019. 100 Questions in Livestock Helminthology Research. Trends Parasitol. 35, 52–71. 10.1016/j.pt.2018.10.006

Morgan, E.R., Lanusse, C., Rinaldi, L., Charlier, J., Vercruysse, J., 2022. Confounding factors affecting faecal egg count reduction as a measure of anthelmintic efficacy. Parasite 29. 10.1051/parasite/2022017

Munguía, B., Saldaña, J., Nieves, M., Melian, M.E., Ferrer, M., Teixeira, R., Porcal, W., Manta, E., Domínguez, L., 2022. Sensitivity of Haemonchus contortus to anthelmintics using different in vitro screening assays: a comparative study. Parasites and Vectors 15, 1–11. 10.1186/s13071-022-05253-3

Osei-Atweneboana, M.Y., Awadzi, K., Attah, S.K., Boakye, D.A., Gyapong, J.O., Prichard, R.K., 2011. Phenotypic evidence of emerging ivermectin resistance in Onchocerca volvulus. PLoS Negl. Trop. Dis. 5, 1–11. 10.1371/journal.pntd.0000998

Page, A.P., Stepek, G., Winter, A.D., Pertab, D., 2014. Enzymology of the nematode cuticle: A potential drug target? Int. J. Parasitol. Drugs Drug Resist. 4, 133–141. 10.1016/j.ijpddr.2014.05.003

Preez, G. Du, Fourie, H., Daneel, M., Miller, H., Höss, S., Ricci, C., Engelbrecht, G., Zouhar, M., Wepener, V., 2020. Oxygen consumption rate of Caenorhabditis elegans as a high-throughput endpoint of toxicity testing using the Seahorse XFe96 Extracellular Flux Analyzer. Sci. Rep. 10, 1–11. 10.1038/s41598-020-61054-7

Prichard, R.K., Geary, T.G., 2019. Perspectives on the utility of moxidectin for the control of parasitic nematodes in the face of developing anthelmintic resistance. Int. J. Parasitol. Drugs Drug Resist. 10, 69–83. 10.1016/j.ijpddr.2019.06.002

Risi, G., Aguilera, E., Ladós, E., Suárez, G., Carrera, I., Álvarez, G., Salinas, G., 2019. Caenorhabditis elegans infrared-based motility assay identified new hits for nematicide drug development. Vet. Sci. 6. 10.3390/VETSCI6010029

Simonetta, S.H., Golombek, D.A., 2007. An automated tracking system for Caenorhabditis elegans locomotor behavior and circadian studies application. J. Neurosci. Methods 161, 273–280. 10.1016/j.jneumeth.2006.11.015

Simonetta, S.H., Migliori, M.L., Romanowski, A., Golombek, D.A., 2009. Timing of locomotor activity circadian rhythms in Caenorhabditis elegans. PLoS One 4. 10.1371/journal.pone.0007571

Suárez, G., Alcántara, I., Salinas, G., 2022. Caenorhabditis elegans as a valuable model for the study of anthelmintic pharmacodynamics and drug-drug interactions: The case of ivermectin and eprinomectin. Front. Pharmacol. 13, 1–9. 10.3389/fphar.2022.984905

Taki, A.C., Byrne, J.J., Wang, T., Sleebs, B.E., Nguyen, N., Hall, R.S., Korhonen, P.K., Chang, B.C.H., Jackson, P., Jabbar, A., Gasser, R.B., 2021. High-Throughput Phenotypic Assay to Screen for Anthelmintic Activity on Haemonchus contortus. Pharmaceuticals 14, 616. 10.3390/ph14070616

Wit, J., Dilks, C.M., Andersen, E.C., Program, B.S., 2022. Nematodes To Understanding Anthelmintic Resistance 37, 240–250. 10.1016/j.pt.2020.11.008.Complementary

